# Relative Principal Components Analysis: Application to Analyzing Biomolecular Conformational Changes

**DOI:** 10.1101/409474

**Authors:** Mazen Ahmad, Volkhard Helms, Olga V. Kalinina, Thomas Lengauer

## Abstract

A new method termed “Relative Principal Components analysis” (RPCA) is introduced that extracts optimal relevant principal components to describe the change between two data samples representing two macroscopic states. The method is widely applicable in data-driven science. Calculating the components is based on a unified physical framework which introduces the objective function, namely the Kullback-Leibler divergence, appropriate for quantifying the change of the macroscopic state as it is effected by the microscopic features. To demonstrate the applicability of RPCA, we analyze the thermodynamically relevant conformational changes of the protein HIV-1 protease upon binding to different drug molecules. In this case, the RPCA method provides a sound thermodynamic foundation for the analysis of the binding process. The relevant collective (global) conformational changes can be reconstructed from the informative latent variables to exhibit both the enhanced and the restricted conformational fluctuations upon ligand association. Moreover, RPCA characterizes the locally relevant conformational changes which can be presented on the structure of the protein.

## 1 Introduction

Studying the transitions and differences between multiple states populated by a dynamic system is a central topic in different fields including chemistry, physics, biology, machine learning and all of data-driven science. A typical task is to uncover how macroscopic changes of the dynamic system are related to the features (variables) that describe its microscopic individuals (instances). Two examples of such microscopic features would be the genetic sequences of a virus taken from snapshots during the course of evolution or the spatial conformations of two biomolecules when they bind to each other. The relationship between the “microscopic” factors of a system and the change of its macroscopic states requires the definition of an appropriate objective function for quantifying the change of the “macroscopic” state of the system. Such a rigorous definition of changes of the macroscopic state of a system in terms of its microscopic features is available for physical systems whose thermodynamic quantities can be measured or computed. For example, the change of free energy (a scalar value) is a suitable quantity to characterize macroscopic changes in physical, chemical and biochemical systems. However, in other areas of data-driven science, such a rigorous definition and quantification of macroscopic changes generally does not exist. Instead, various heuristic objective functions are used in practice. Examples include divergence measures from information theory (Lin, 1991) and the wide variety of objective functions which are used for prediction and feature extraction in pattern recognition (Fukunaga, 1990). Mining the factors informative for the change between two samples is of high importance and of general interest in all areas of data-driven science and is generally performed in a high-dimensional feature space. In fact, mining informative features is the central theme in a large domain of machine learning and includes methods such as dimensionality reduction (Murphy, 2012; Theodoridis, 2015), feature extraction (Fukunaga, 1990), and latent variable models (Bishop, 1998). However, one needs to select an objective function that is appropriate for quantifying the change before applying a multivariate method to extract the informative features.

Analyzing the conformational changes taking place during biomolecular reactions is one of the most important tasks in structural biology. Unfortunately, analyzing and mechanistically understanding biochemical interactions is quite tricky due to the complex conformational dynamics in the high-dimensional space where the interactions take place. The macroscopic changes in biochemical systems, on the other hand, are quantified using the change of free energy (a scalar quantity). Molecular simulations are becoming a more and more attractive tool for analyzing conformational changes of biomolecules. Current interest in analyzing molecular conformational changes (e.g. using Markov state models (Husic and Pande, 2018)) focusses on characterizing the kinetic changes between representative conformations within one macroscopic state and the analysis is performed in a data space where the points are the (clustered) conformations. Methods such as PCA (Amadei et al., 1993; Kitao and Go, 1999) and partial least squares (Krivobokova et al., 2012) were used to analyze the conformational changes in the feature space (coordinates). However, most of these methods are only suited for studying the dynamic changes within one macroscopic state and not for studying transitions and changes between two states. On the other hand, methods adapted from multivariate analysis often lack thermodynamic insight.

In this work, we introduce a unified framework rooted in statistical information theory and statistical mechanics (Kullback, 1997a; Barndorff-Nielsen, 1978; Ahmad et al., 2015a, 2017) for the purpose of studying the change between two data sets representing two states. A new method termed *“Relative Principal Components Analysis*” is introduced to extract the optimal relevant principal components describing the change between two data samples (two states). This method is applicable in all areas of data-driven science and is introduced in its generality in Section 2 and 3. As special but important example, starting in Section 4, we apply RPCA for analyzing the energetically relevant conformational changes of a biomolecule (protease of the human immunodeficiency virus, HIV-1) upon binding to various ligands.

## 2 A general formalism for analyzing changes in dynamic systems based on information theory

Before going into the technical details of finding the directions in feature space that are informative of the change between two states, we first introduce a physical framework for defining and quantifying the change of dynamic systems in all areas of data-driven sciences and justify the objective function used for quantifying the macroscopic change.

Let ***x*** be a random vector encoding the microscopic features (variables) which are necessary to identify the difference between the instances of a system of interest. By a *system* we mean a collection of individuals or *instances,* e.g. viruses or molecules, where each instance is defined via a set of (microscopic) *features*. A macroscopic state (*a*) of the system is defined by the probability density distribution *P*(*x|a*) = *P_a_*(*x*) of all possible instances (***x***) when the system is at equilibrium in this macroscopic state *a.* The macroscopic state *a* can be changed into a new macroscopic state *b* by perturbing the probability distribution of the microscopic instances ***x*** to *P*(***x***|*b*) = *P_b_*(***x***) Here, a Bayesian approach is used whereby the macroscopic state is viewed as a random variable, and the conditional probability *P*(***x***|*i*) = *P_i_*(***x***) is naturally interpreted as the equilibrium distribution of state *i* that can be taken from a finite or discrete set. A relationship between the two macroscopic states can be obtained by applying Bayes’ theorem, yielding the following equation for the probability density ratio *P_b_*(***x***)/*P_a_*(***x***):

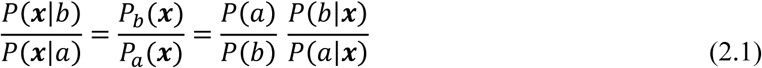

Here, *P*(*b*)and *P*(*a*) are conditional probabilities under the (implicit) condition of the perturbing factors. Averaging the logarithm of the density ratio equation over the probability distribution of state *b* (*P_b_*(***x***)) yields:

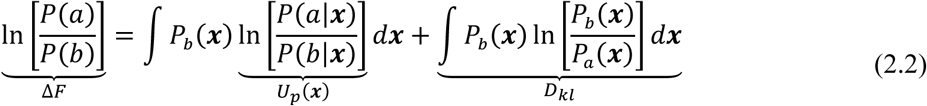

Indeed, this is a general derivation of the Perturbation Divergence Formalism (PDF), which was previously derived for physical systems for the purpose of decomposing the change of free energy (∆*F*) between two macroscopic states *a* and *b* (Ahmad et al., 2017, 2015a):

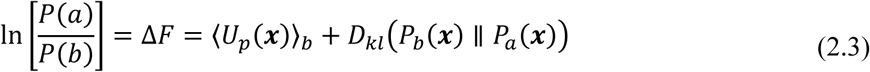

For physical systems, we use here the natural unit of the energy (*kT*)^−1^ = 1. *D_kl_* is the Kullback-Leibler (KL) divergence (Kullback and Leibler, 1951) (also termed the relative entropy (Cover and Thomas, 2006) or the discriminant information (Kullback, 1997a)) between the probability distributions of states *a* and *b*. Interestingly, equation (2.2) provides a purely probabilistic formalism of free energy change via a new probabilistic definition of the perturbation *Up* of the microscopic instances as the logarithm of the ratio of the posterior probabilities of two macroscopic states given a particular microscopic configuration ***x***:

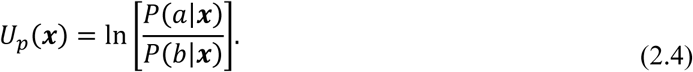

### Bridging the macroscopic change and the microscopic features

The concept of the perturbation of a microscopic instance was originally introduced in statistical mechanics as an energetic quantity in order to formalize the relationship between the microscopic (atomistic) description of a physical system and its macroscopic changes between states (free energy change)(Kirkwood, 1934; Zwanzig, 1954). When considering the change between two macroscopic states, the perturbation *Up*(***x***) is a one-dimensional (microscopic) variable which has the same KL divergence (discriminant information) as the high dimensional feature ***x*** (Ahmad et al., 2015a):

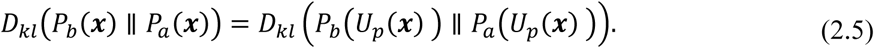

In statistical inference theory (Berger and Casella, 2001; Lehmann and Casella, 2003), such a variable is termed a sufficient statistic. A sound framework for the relationship between the change of macroscopic states of a dynamic system and its microscopic elements can be inherited from the formalism of exponential families (Barndorff-Nielsen, 1978; Brown, 1986) in statistical estimation theory.

Let us assume we have a dynamic system at equilibrium in a macroscopic reference state labeled by the so-called *natural parameters λ.* The system can populate a new macroscopic state *P_λ_*(***x***) upon perturbing its original state 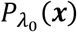. The principle of minimum discrimination information (Kullback, 1997b) (equivalent to the principle of maximum entropy (Greven et al., 2003)) is applied to find the new distribution *P_λ_*(***x***) via minimizing the KL divergence from the reference distribution 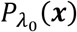 to the new distribution *P_λ_*(***x***) under the constraint of a finite expected value of the sufficient statistic 〈***T***(***x***) 〉_*λ*_. The probability density distribution *P_λ_*(***x***) forms an exponential family of distributions in terms of the reference distribution:

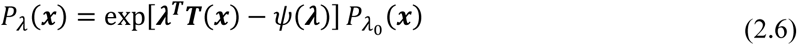

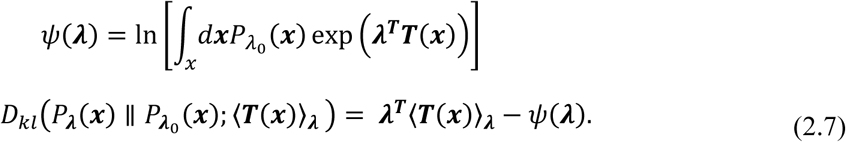

Here, the sufficient statistic ***T***(***x***) is a vector (or a scalar) function of the microscopic configuration ***x*** of the system. The cumulant generating function *ψ*(***λ***) depends only on the natural parameters ***λ***. Besides being used to formulate a theoretical framework for studying the error bound of parameter estimation (Kullback, 1997b), the formalism of exponential families plays a central role in different fields of machine learning such as generalized linear models and variational inference (Murphy, 2012). In statistical thermodynamics, Kirkwood introduced his thermodynamic integration (TI) equation (Kirkwood, 1934) using the exponential family to “*alchemically”* interpolate two macroscopic states (Kirkwood, 1934). Indeed, the one-dimensional sufficient statistic, “the perturbation”, is the appropriate tool for interpolating between two macroscopic states in free energy calculations. Generally, a higher-dimensional sufficient statistic is required when studying multiple macroscopic states (Berger and Casella, 2001). The sufficient statistic and the corresponding cumulant generating function in equations (2.6) are not unique. Notably, comparing equations (2.2) and (2.7) shows that the sufficient statistic can be selected such that the corresponding cumulant generating function is a function of the change of free energy between the macroscopic states of dynamical systems. For example, when selecting the negative value of the perturbation (–*U_p_*(***x***)) as a sufficient statistic for studying the change between two macroscopic states, the corresponding cumulant generating function is the negative of the free energy change and equation (2.6) turns into the free energy perturbation equation that is well known in statistical thermodynamics (Zwanzig, 1954). Another example is the logarithm of the density ratio, which is a well-known sufficient statistic in statistical inference, and its corresponding cumulant generating function is the relative change of free energy (see Supplementary Material S4). An interesting known relationship of an exponential family is the functional dependence of the change of *ψ*(***λ***) (the function of the macroscopic state) on the microscopic features ***x*** that is given by the average and the covariance of the sufficient statistic:

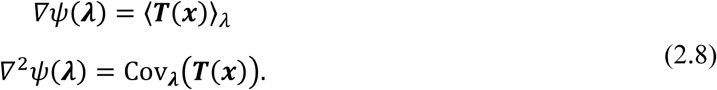

The free energy thermodynamic integration equation (Kirkwood, 1934) is a special case of equation (2.8). However, quantification of the macroscopic change is not of general interest in the field of data-driven sciences. The important task here is identifying the unknown sufficient statistic that explains the influence of the microscopic features of the macroscopic change. Unlike free energy change, KL divergence is a quantity familiar in machine learning and can be computed using parametric or nonparametric models (Sugiyama et al., 2012). Indeed, equation (2.7) shows that KL divergence is a Legendre transformation (Barndorff-Nielsen, 1978; Rockafellar, 1970) of the cumulant generating function and can be used to quantify the change of the macroscopic state. The PDF given by equation (2.2) is a special case of equation (2.7) where we use the perturbation as a sufficient statistic for studying the change between two macroscopic states (*λ* = 0,1). Due to the Legendre duality (Barndorff-Nielsen, 1978; Rockafellar, 1970) between the KL divergence and the change of the free energy, the relevance of the PDF goes beyond being a decomposition of the change of free energy. In fact, the terms of equation (2.2) are the perturbations (the features informative of the change) and their fingerprints (the configurational changes) which are quantified by the KL divergence. In this view, significant perturbations are reflected by significant changes of KL divergence.

## 3 Relative principal components analysis

This section presents a new method for analyzing the change between two states (data sets) using multivariate analysis methods (Theodoridis, 2015). Studying the change between multiple states is beyond the scope of this work and will be presented in a future publication. The newly introduced method termed “relative principal components analysis” (RPCA) computes collective canonical variables (linear combinations of the original features) termed the relative principle components (RPCAs) to which the KL divergence factorizes additively. Indeed, factorization of the KL divergence is equivalent to factorization of the logarithm of the logarithm of the density ratio which is a sufficient statistic of interest in machine learning (Sugiyama et al., 2012) (see above). Factorization of KL divergence was introduced for multivariate normal distributions in the seminal work of Kullback (Kullback, 1997a). However, the theoretical approach of factorizing KL divergence as introduced by Kullback was not accessible in practice. Specifically, a solution is needed around the singularity of the covariance matrices and the resulting features have to be optimal with respect to maximizing their KL divergences (see below).

Let ***x*** = (*x*_1_ … *x_d_*)^*T*^ be a *d*-dimensional continuous random variable with two samples from two macroscopic states that will be labeled (*a*) and (*b*). RPCA aims at finding *k* latent canonical variables ***y*** = (*y*_1_,*y*_2_ … *y_k_*)^*T*^ = *f*(***x***) which satisfy the following two conditions: (i) their marginal distributions are independent in the states *a* and *b* meaning that their KL divergences are additive, and (ii) optimal in terms of maximizing the KL divergence, such that we can use *m* (*m* ≪ *d*) latent variables to represent the significant directions informative of the change. The KL divergence of a new variable ***y*** = *f*(***x***) is always non-negative (Kullback, 1997a; Cover and Thomas, 2006) and is bounded from above by the KL divergence of the original variable ***x*** (Ahmad et al., 2017; Kullback, 1997a):

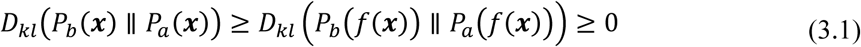

The new variable ***y*** = *f*(***x***) is “sufficient” if equality holds in (3.1). When studying the change between two macroscopic states, a sufficient one-dimensional variable always exists (e.g. the perturbation (Ahmad et al., 2017)) regardless of the dimensionality of ***x***. Although the existence of a one-dimensional sufficient feature appears promising, it is not practically useful for two reasons: (i) the analytical nature of the sufficient one-dimensional variable is generally unknown, and (ii) the complexity and nonlinearity of a sufficient one-dimensional variable –if it is known– will hinder its interpretability in terms of the original features ***x*** In fact, its simple interpretability and analytical traceability is one reason for the wide-spread use of the Gaussian linear parametric model in latent variable models (Bishop, 1998) (e.g. PCA).

We will adopt this model here as well. Then the latent variables are linear combinations of the original variable and are normally distributed:

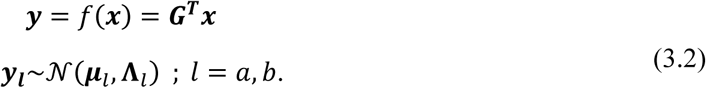

Here, ***G*** is a *d* × *k* transformation matrix with columns ***g_i_***. Thus 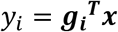 and ***μ***_*l*_; *l* = *a,b* are the averages of the two distributions. The covariance matrices ***S***_*l*_ of the original variables are related to the covariance matrices of the latent variables by **Λ**_*l*_ = ***G^T^S***_*l*_***G***; *l* = *a*, *b*. Under the model assumption of normality, the independence of the variables ***y_i_***requires **Λ**_*l*_ to be diagonal in both states *a* and *b*:

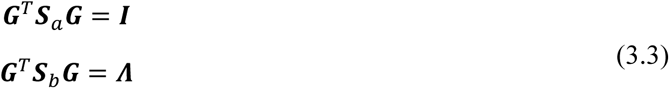

Here, the diagonal matrix of state *b,* **Λ** = diag(*λ*_1_ … *λ_k_*), contains the variances *λ*_1_ … *λ_k_* of the latent variables at state *b* while the covariance matrix of the latent variables at state *a* is arbitrarily selected to be an identity matrix (***I***). In case of a multivariate normal distribution (***S***_*a*_ is nonsingular), Kullback (Kullback, 1997a) formulated the solution for ***G*** as the generalized eigenvectors corresponding to the generalized eigenvalue problem |*S_b_* – *λ****S***_*a*_| which can be solved using Wilkinson’s algorithm (Martin and Wilkinson, 1971) involving the Cholesky decomposition of ***S***_*a*_ (Stewart, 2001, p. 229). Practically, the covariance matrices of real data are mostly singular or ill-conditioned and the generalized eigenproblem is not solvable. Singularity arises due to the fact that the real dimensionality of the probability distributions is smaller than the apparent dimensionality of ***x*** (Mardia et al., 1979, p. 41).

### RPCA via simultaneous diagonalization of two matrices

Here, we present a general algorithm for the simultaneous diagonalization of the matrices in equation (3.3) which can be used even if ***S***_*a*_ is singular.

A transformation matrix G ∈ ℝ^*d*×*k*^ that simultaneously diagonalizes the symmetric matrices ***S***_*a*_ ∈ ℝ^*d*×*d*^ (of rank *k*) and ***S***_*b*_ ∈ ℝ^*d*×*d*^ (equation (3.3)) can be found by a combination of two transformation matrices:

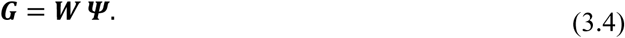

1) The so-called *whitening transformation* (Fukunaga, 1990) matrix G ***W*** ∈ ℝ^*d*×*k*^ of the matrix ***S***_*a*_ is computed from its eigendecomposition 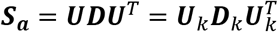:

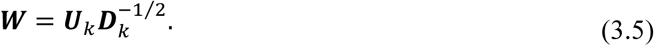

Here, the *k* eigenvectors in the columns of ***U***_*k*_ correspond to the *k* nonzero eigenvalues in the diagonal matrix ***D***_*k*_ Clearly, ***W*** reduces ***S***_*a*_ to an identity matrix ***W^T^ S***_*a*_***W*** = ***I*** ∈ ℝ^*k*×*k*^ corresponding to the covariance matrix of the whitened data (***W^T^x***). The algorithm above is well known in case ***S***_*a*_ is nonsingular (*d* = *k*), (Fukunaga, 1990).

2) The matrix ***Ψ*** is formed using the eigenvectors from the symmetric matrix ***W^T^S***_*b*_***W***:

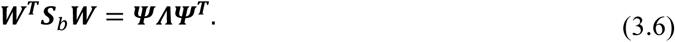

It is straightforward to see from (3.5) and (3.6) that ***G*** = ***W Ψ*** simultaneously diagonalizes ***S***_*a*_ and ***S****b* and satisfies equation (3.3).

The *k* columns of ***G*** are the generalized eigenvectors with the corresponding generalized eigenvalues in the diagonal matrix ***Λ***. The relative principle components 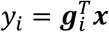 can be reordered based on their KL divergences. The additive KL divergences of the independent variables *y_i_* can be analytically computed based on the model assumption of normality (Kullback, 1997a):

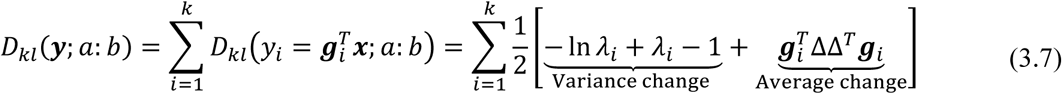

Here ∆= ***μ***_*p*_ – ***μ***_*a*_ is the change of the average of the distributions of ***x*** in states *a* and *b*. However, it is important to keep in mind that the value of *D*_KL_ of the components from equation (3.7) is computed based on the model assumption (normality). Fortunately, a significant KL divergence of a variable *y_i_* has to be reflected in a significant change between its distributions in states *a* and *b* which can be used to assess the accuracy of the model-based KL divergences (see the example below).

### Optimal RPCAs via average-covariance sub-spacing

Although the simultaneous diagonalization algorithm above returns independent latent variables into which the KL divergence factorizes additively, the obtained latent variables are not optimal in terms of maximizing the KL divergence. Optimal latent variables are required for reconstructing the most informative approximation (maximizing KL divergence) of the original variable; see below. The KL divergences of the relative principle components in equation (3.7) can be decomposed into the terms holding the change of the variances 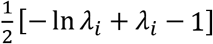 and the terms holding the change of the average 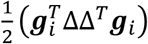. Indeed, the latent variables from equation (3.4) are optimal with respect to maximizing the KL divergences due to change of the variances (Kullback, 1997a). Unfortunately, this optimality is violated by the contributions to the KL divergences due to the change of the average 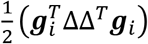. Therefore, the following average-covariance sub-spacing algorithm is introduced to achieve the optimality of RPCAs with respect to maximizing their KL divergences. The detailed derivation is presented in Supplementary Material S1. The idea is to find a transformation matrix ***G*** = [***g***_*μ*_ ***G***_*υ*_] such that the KL divergence of the variable 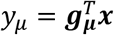 summarizes the total KL divergence due to changes of the averages while the KL divergences of the variables 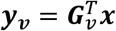 are purely due to the change of covariance. The required solution for ***g***_*μ*_ is given by:

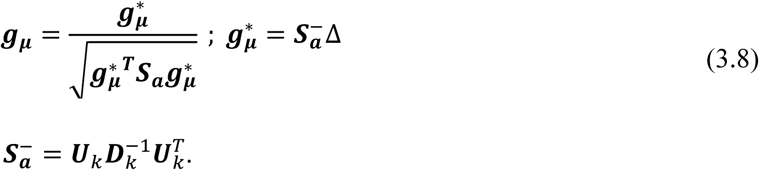

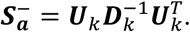

Here, the generalized pseudoinverse 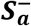 is used for the general case (e.g. singular ***S***_*a*_) and ***g***_*μ*_ is normalized with respect to the covariance matrix ***S***_*a*_ such that 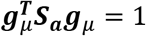. The KL divergence of the variable 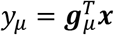 includes the total KL divergence due to the change of the average, which in turn equals half of the Mahalanobis distance between the averages 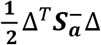:

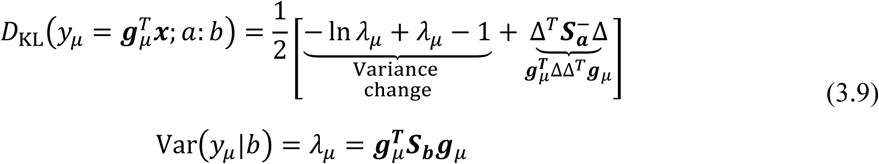

Here, the KL divergence of the new variable 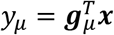 also includes a contribution due to the change of its variance (*λ_μ_*).

The remaining generalized eigenvectors ***G***_*υ*_, which do not contain contributions arising from the change of the average 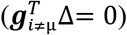, are computed after deflating the contributions to the KL divergence due to vector ***g***_*μ*_ from the matrices ***S***_*a*_ and ***S***_*b*_ Given the vector ***g***_*μ*_ the restricted simultaneous diagonalization problem can be presented in the equations:

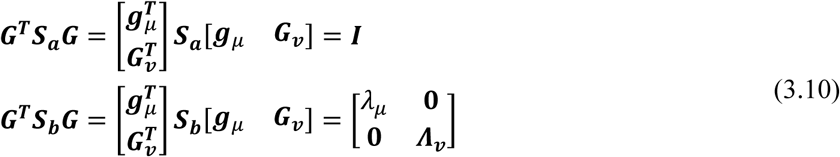

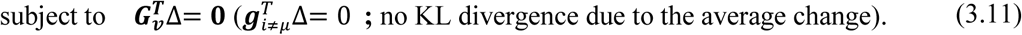

Here, ***Λ***_*υ*_ is a diagonal matrix and **0** denotes a matrix or a vector of zeros of suitable dimensionality. To fulfill the conditions in (3.10) and (3.11), the generalized eigenvectors ***G***_*υ*_ can be constructed using a combination of two transformation matrices (for details see Supplementary Material S1):

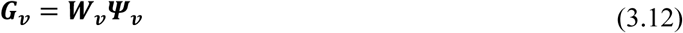

(i) The whitening transformation matrix ***W****υ*, similarly to equation (3.5), is constructed here from the eigendecomposition of the covariance matrix of the projection of ***x*** on the subspace, which is orthogonal to the vector (***S***_*a*_***g***_*μ*_). (ii) The second transformation *Ψ_υ_* is obtained from the eigenvectors of covariance matrix of the projection of the whitened data (***W^T^x***) onto the subspace which is orthogonal to the vector 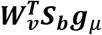. Here, the number of generalized eigenvectors from the optimal RPCA sub-spacing is one less than the number of generalized eigenvectors from the non-optimal RPCA algorithm of equation (3.4).

### Data reconstruction from the latent variable

The influence of the *m*-dimensional (*m* < *d*) latent variable 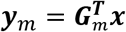 on the original *d*-dimensional variable x ∈ ℝ^*d*^ can be presented via the reconstruction (projection) *x̂* ∈ ℝ^*d*^ in the subspace which is spanned by the corresponding *m* generalized eigenvectors ***G***_*m*_ = [***g***_1_ *…* ***g***_*m*_] and is given by the relationship (Harville, 2008):

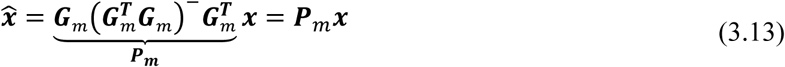

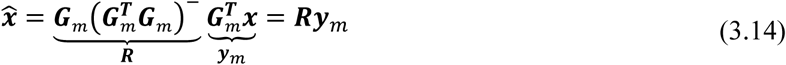

Here, ***P***_*m*_ ∈ ℝ^*d*×*d*^ is the projection matrix (Harville, 2008) on the *d*-dimensional subspace which is spanned by columns of ***G***_*m*_ ∈ ℝ^*d*×*m*^ is the reconstruction matrix facilitating the transformation from the latent variable. It is straightforward (see Supplementary Material S2) to show that the KL divergence between the distributions of the reconstructed variable *x̂* equals the KL divergence between the distributions of its corresponding latent variable ***y***_*m*_. In other words, the most informative approximation (the one maximizing KL divergence) of the original variable ***x*** using a restricted number (*m* < *d*) of variables is obtained using the *m*-dimensional latent variables ***y***_*m*_ with the highest KL divergences.

### Change hotspots from RPCA

Besides representing the changes collectively (incorporating all *x_i_*), RPCA provides the possibility to map the feature-wise (local) contributions to the change from the individual elements *x_i_* of ***x*** such that the elements of ***x*** with larger contribution to the change (KL divergence) can be interpreted as the hotspots of the change. The contributions to the divergence in equation (3.7) can be approximated as a sum of two quadratic terms:

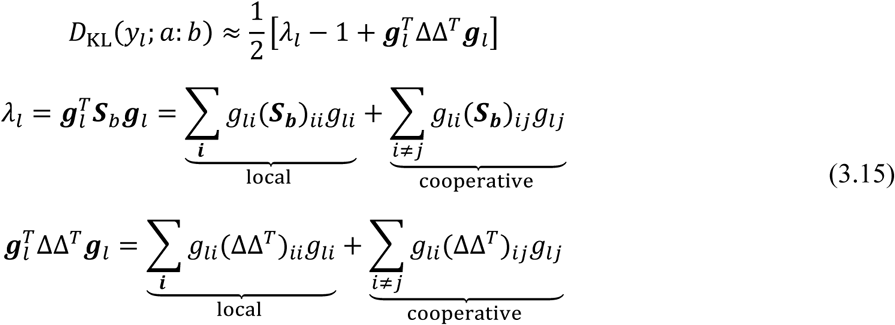

Here, (***A***)_*ij*_ denotes element (*i,j*) of matrix ***A***. The contributions to the quadratic terms are due to the local contributions from the elements and their cooperative (cross) terms taking into account that each element of a generalized eigenvector (*g_li_*) corresponds to an element *x_i_*. However, it is clear that the contributions to the quadratic terms in equation (3.15) can be collected via arbitrary grouping of the elements into subgroups. The computation of the (local) group-wise contributions and their cooperative contributions is equivalent to the computation of the quadratic terms using the corresponding submatrices of ***S***_*b*_ and ∆∆^*T*^.

### Asymmetric nature of RPCA

Multivariate analysis methods can be grouped into methods handling either one state or multiple states. The methods which study one state include methods handling one multivariate variable, such as principal component analysis, factor analysis and independent components analysis, and methods handling two multivariate variables with concurrent measurement (a joint distribution) such as canonical correlation, regression and partial least squares. Discriminant analysis (classification) methods, on the other hand, aim at finding the variation between several states (classes). Although RPCA is similar to feature extraction for classification (Fukunaga, 1990), there are fundamental differences between pattern classification and the physical definition of the change, which is introduced above. For example, the discriminant features in Fisher discriminant analysis (FDA) are related to the change of the averages of distributions and the information from the covariance matrices is whitened using a unified pooled (average) covariance matrix (Fukunaga, 1990). Therefore, the usage of FDA for dimensionality reduction (Sugiyama et al., 2010) is known to be restricted by a limit on the number of dimensions which is equal to the number of classes minus one. While the objective functions in discriminant analysis are required to be symmetric (Fukunaga, 1990), the objective function (KL divergence) for RPCA is asymmetric, which reflects the directed character of changes (Feng and Crooks, 2008). When studying the backward change from state *b* to state *a,* the generalized eigenvectors ***G***_*R*_ pertaining to the reverse direction are the scaled eigenvectors pertaining to the forward direction (***G***) and the generalized eigenvalues, which present the change of the variance, are inverted (1/*λ_i_*):

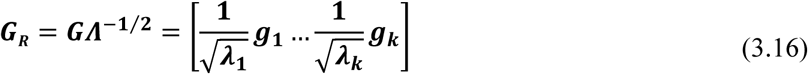

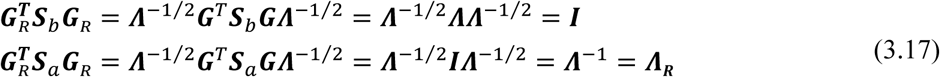

Taking into account that the KL divergence due to the change of the variance 1/2 (− ln *λ_i_* + *λ_i_* – 1) in equation (3.7) is a convex function with a minimum of 0 at *λ_i_* = 1, we can divide the relative principal components into two groups. The first group includes the components with *λ_i_* > 1 where the variance (quantified by *λ_i_*) along the direction ***g***_*i*_ increases as we transit from state *a* to *b*. The second group includes the components with *λ_i_* < 1 since the respective variance decreases as we transit from state *a* to *b.* When the backward change is taking place from *b* to *a,* the roles of the components of these groups (quantified by 1/ *λ_i_*) are exchanged. The same role of these groups is also observed when taking the KL divergence due to the change of the average where the directions ***g***_*i*_ with *λ_i_* > 1 have diminished contributions 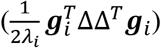 to the backward direction from *b* to *a* in comparison with their contribution 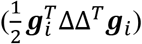 to the transition from state *a* to *b* and vice versa.

### RPCA and the distance metric

Even though the task of RPCA (analyzing the change between states) differs from the task of PCA (finding the variation within one state; see Figure 1), a similarity exists regarding the applied distance metric. However, it is important to notice that the generalized eigenvectors are orthonormal with respect to the matrix (Harville, 2008) 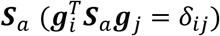, orthogonal with respect to 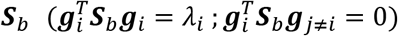 and not necessarily orthonormal to each other 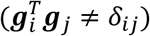. RPCA analysis can be interpreted as using the distance metric (Dryden and Mardia, 2016) of state *a* to analyze state *b.* Indeed, applying the whitening transformation (Fukunaga, 1990, p. 31) in equation (3.5) removes the “information” within state *a* (***W^T^S***_*a*_***W*** = ***I***). The eigendecomposition of ***W^T^S***_*a*_***W*** in (3.6) is performed in the whitened space in which the squared Euclidean distance between two points ***x***_1_ and ***x***_2_ equals their Mahalanobis distance in the original space and the projections on the generalized eigenvectors factorize the Mahalanobis distance (see Supplementary Material S3):

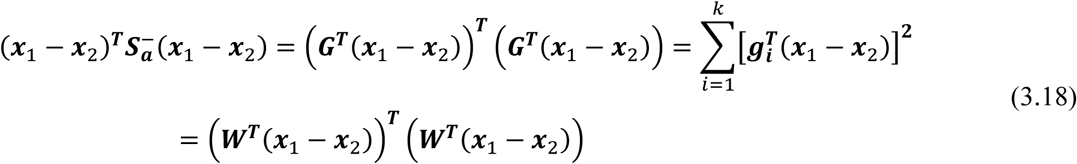

**Figure 1.**
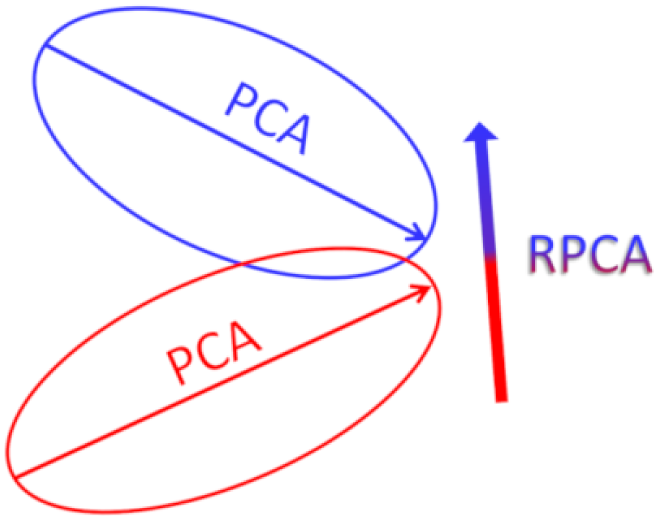
Schematic representation of the conceptual difference between PCA which finds the largest variation within each state and RPCA which finds the change from the initial to the final state.

Moreover, the average Mahalanobis distance of points in state *b* to the average of state *a* can be written as the sum of the generalized eigenvalues and the Mahalanobis distance between the averages of the states (see supplementary material S3):

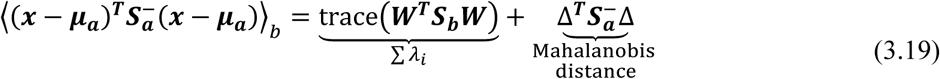

## 4 Application of RPCA to reveal the thermodynamically important molecular conformational changes

Although our RPCA method is not limited to biomolecular data, the starting point for its development was based on the new insight stemming from our recently introduced perturbation-divergence formalism (PDF) (Ahmad et al., 2017, 2015a). PDF reveals that the KL divergence (*D_kl_*) equals the work which is spent to change the conformational ensemble and hereby is the thermodynamically relevant objective function for analyzing conformational changes (Ahmad et al., 2015a). Indeed, PDF provides a flexible framework for studying conformational changes involving either all atoms of the structure, a part of them, or aspects of the structure based on phenotypical features (e.g. open/closed, pocket volume, domain movement, etc.); see the discussion in (Ahmad et al., 2017). An important benefit of using the KL divergence for quantifying the energetics related to conformational changes is avoiding the misleading entropic terms which are subject to enthalpy-entropy compensation when using the changes in conformational entropy to estimate the importance of conformational changes, as was previously shown (Ahmad et al., 2016, 2015b, 2017).

In the following, we present use cases for applying RPCA to analyze the molecular conformational changes of the protein HIV-1 protease upon binding to several inhibitor molecules. The conformations of the protein are sampled via molecular dynamics simulations for both the initial (free, unbound) state and the final (bound) state. Technical details on these simulations are provided in Supplementary Material S5. Figure 2 shows an assessment of the relationship between the relevant components of the dynamic change within one state, which are analyzed using traditional PCA of the data points of the final state, and the thermodynamic importance of the corresponding conformational changes. The thermodynamic importance of the conformational changes along a principal component to the association process is quantified by the KL divergence of the distributions of projections of both the free and the bound state conformations on the component (Ahmad et al., 2016). Figure 2a shows that the eigenvalues of the principal components are not related to their thermodynamic importance for the association process. To illustrate this more clearly, Figure 2b shows that the principal component 3654 has more thermodynamic importance (KL divergence) than the first principal component, which is mostly irrelevant for the association process.

**Figure 2.**
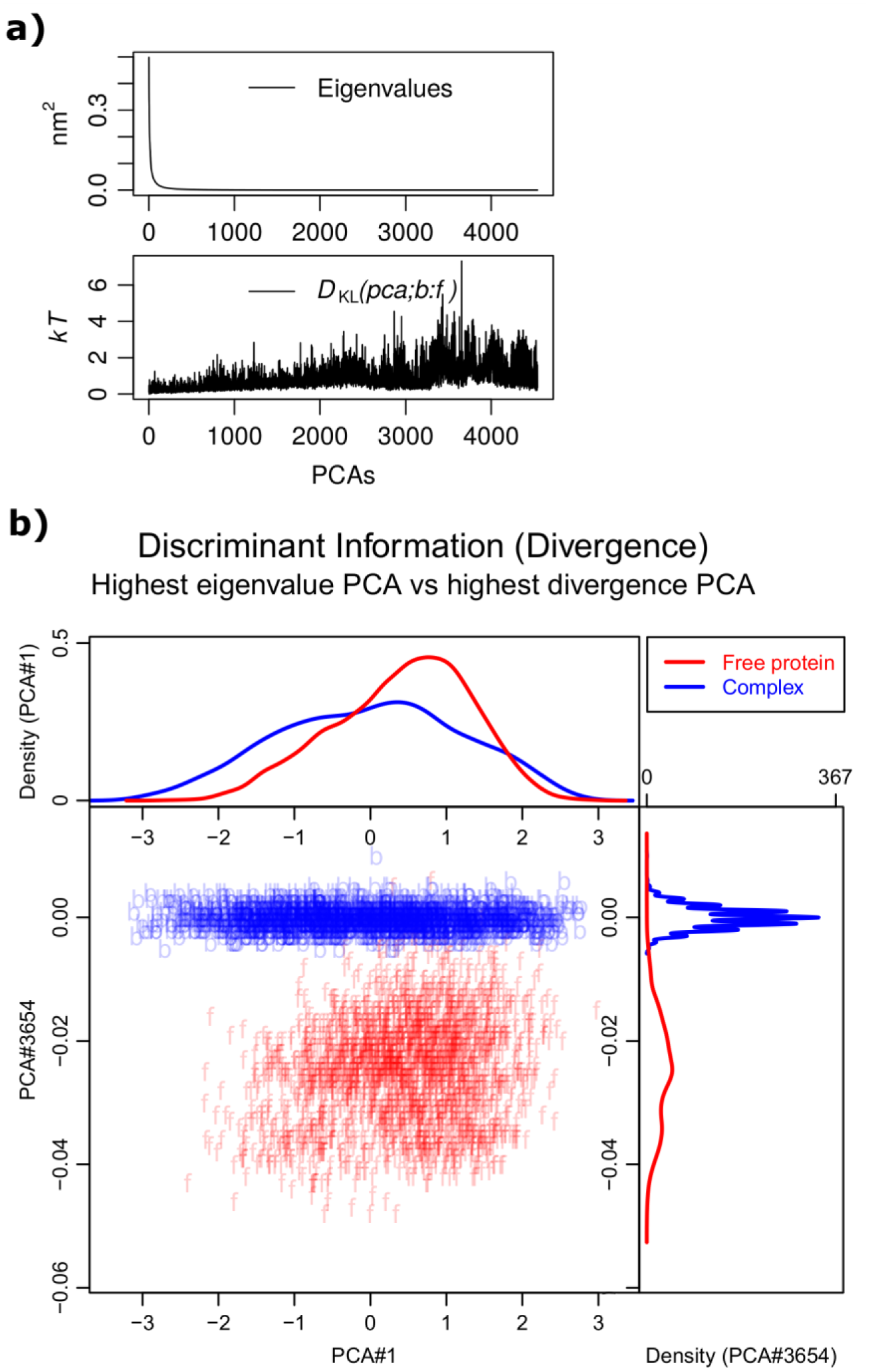
Assessing the thermodynamic contribution of the conformational changes along the directions of the principal components from PCA to the association process. **a)** Top panel: Shown are PCA eigenvalues derived from the covariance matrix of the conformational fluctuation (heavy atoms) in an MD simulation of HIV-1 protease bound to its inhibitor Tipranavir. Bottom panel: Shown is the importance (KL divergence) along PCAs computed from the projections of the data points (conformations) of the final state (bound) and the initial state (free). b) The divergence of the largest eigenvalue PCA#1 (*D_kl_* ≈ 0.1) is compared to that of the PCA with highest divergence #3654 (*D_kl_* ≈ 7.3). 2000 data points are projected on the PCA vectors of the final (bound; blue) and the initial (free; red) states. The marginal probability distribution densities are shown on the side panels. The PCA analysis is performed on the conformations from the bound state using the heavy atoms of the protein. The superposition of the conformations of both the bound and the free state ensembles and the superposition of the two ensembles to each other are performed in a way similar to RPCA (see below). The KL divergences between the distributions of the projections of the two states are computed using the KLIEP method (Sugiyama et al., 2008).

Figure 3 shows the RPCA of the conformational changes of wild-type HIV-1 protease upon binding to a drug molecule that has high affinity (Tipranavir). Presented are both the non-optimal RPCA (Figure 3a) and the optimal RPCA calculated with the sub-spacing method (Figure 3b). The scores (KL divergence) of the importance of the components show that the first few components account for most of the KL divergence. The data points (conformations) of both the free, unbound state and the bound state are projected on selected components to illustrate the correspondence between the score (model based) and the real divergence which can be extracted from the difference between the distributions of the projections of the states. There are clear benefits of the optimal sub-spacing RPCA (Figure 3b). Namely, the first component has a significant divergence because it collects the total contribution to the KL divergence arising from the change of the average. In contrast, one obtains identical averages when projecting the conformations sampled in the two states onto the remaining components. Thus, the remaining components do not contribute to the KL divergence that is due to the change of the averages. Figure 4 shows a comparison of the KL divergence (discriminant information) of the top-scoring component versus the lowest-scoring and the principal component with largest eigenvalue from PCA of the bound state.

**Figure 3.**
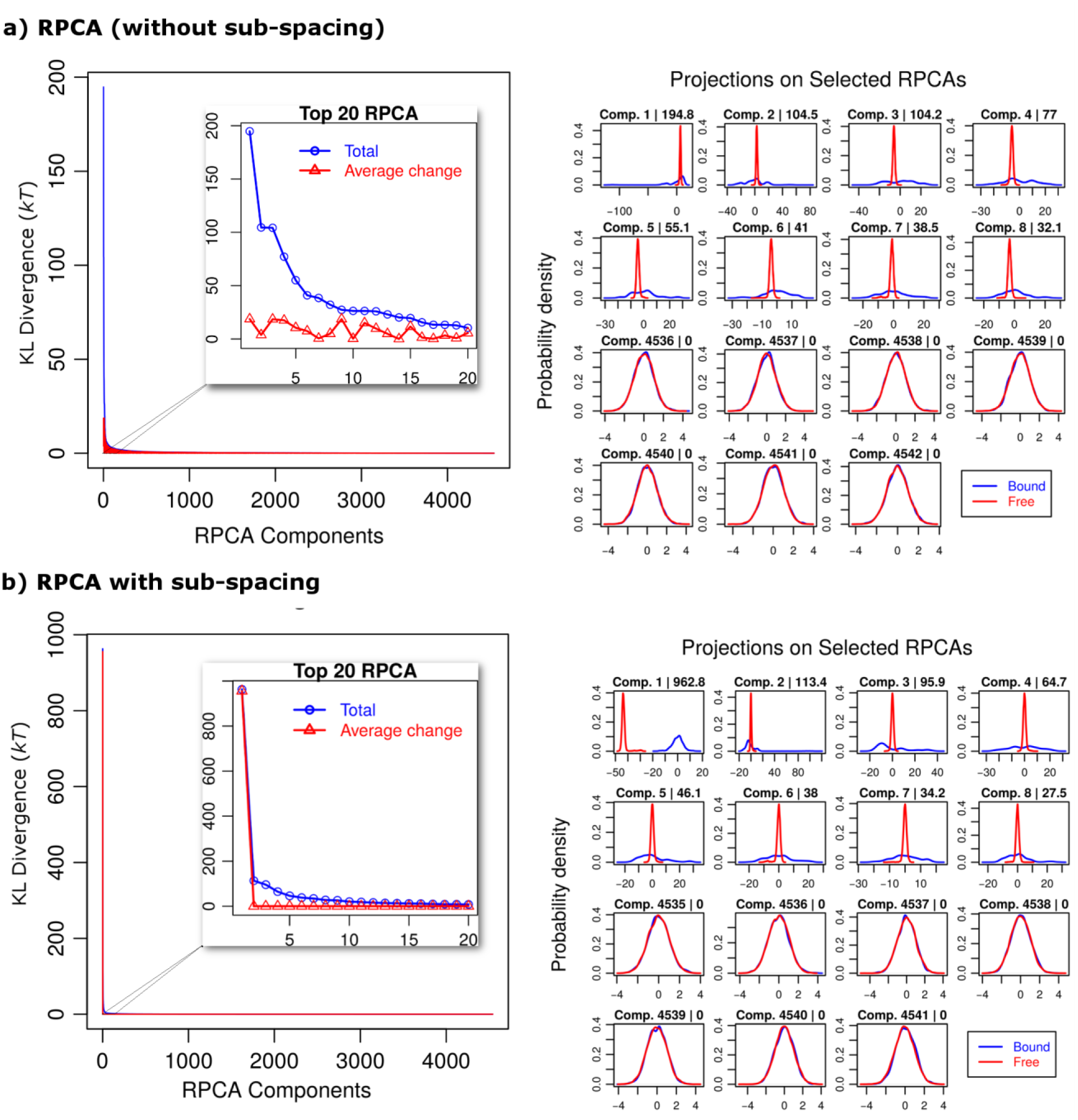
RPCA of the conformational changes of HIV-1 protease upon its association with Tipranavir. Shown are the non-optimal RPCA (a) from equation (3.4) and the optimal RPCA with sub-spacing (b) from equation (3.12). The right panels show the KL divergences of the components (blue colored) and their corresponding contributions due to the change of the average (red colored). Left panels show projections of the data points (conformations) of both the initial (free, unbound) and the final bound state on selected components. The scores (KL divergences) of the components are displayed after their corresponding number. The projections show that the components (components 1-8) with the highest rank (KL divergence) distinguish the change between the free and bound states while the components with the lowest rank do not distinguish the change (similar projections). The analysis is performed using the heavy atoms of the protein. The plots are generated using R (R, 2012) and the densities are smoothed using the kernel density estimation.

**Figure 4.**
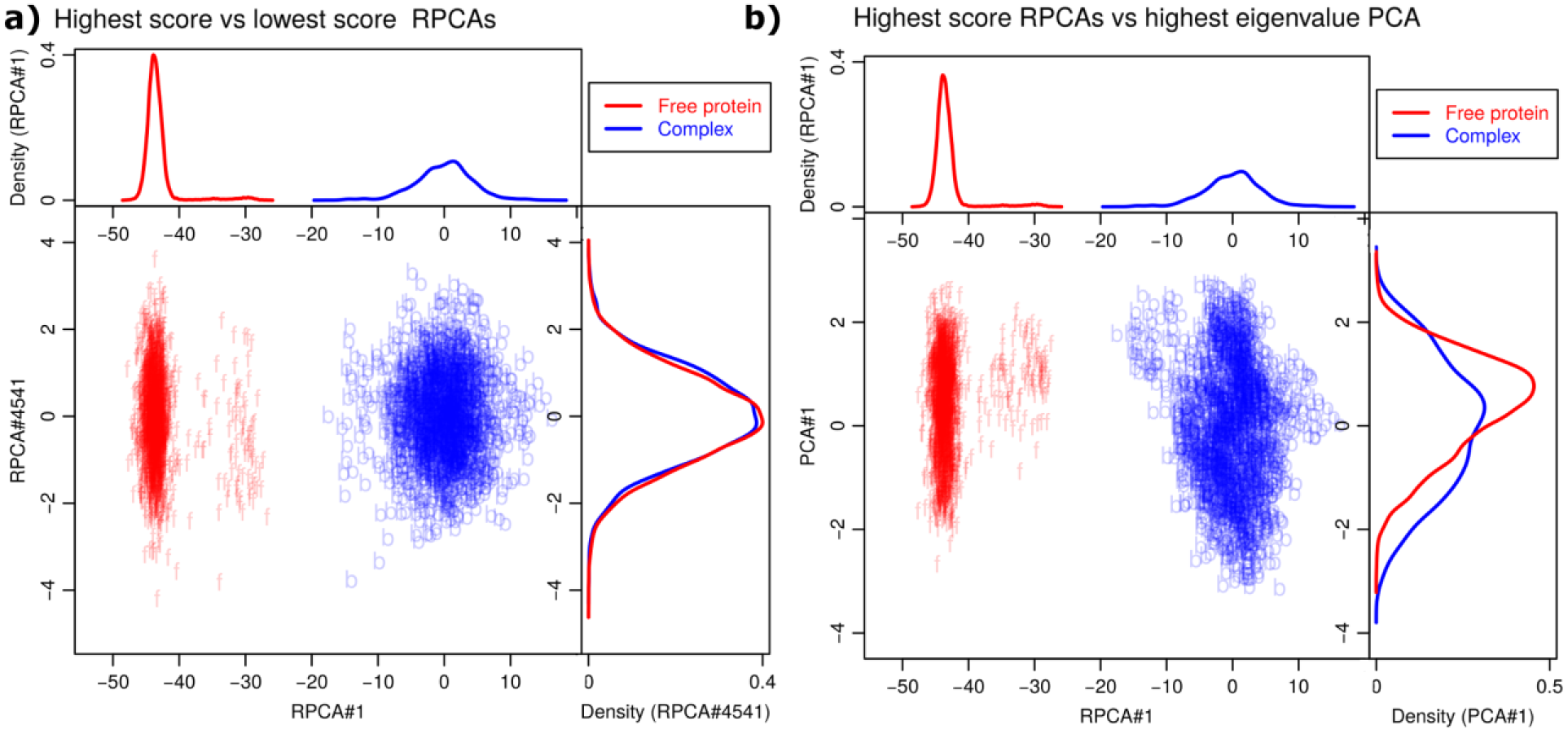
KL divergences of the highest scored relative principal component. The initial state (red) and the bound state (blue) are mapped on the top-scoring RPCA#1 against the lowest-scoring RPCA#4541 (a) and the top-scoring RPCA#1 against the top-scoring PCA#1 (highest eigenvalue) (b). RPCA is successful in extracting the components holding the most significant information on the change (binding process from free to bound state). The marginal probability distribution densities are shown on the side panels. Samples of 2000 data points where used for the projections.

RPCA facilitates studying the conformational changes from both the collective and the local point of view. The relevant collective conformational changes (with respect to their KL divergence) can be presented via reconstructing the conformations in the subspace, which is spanned by the relevant generalized eigenvectors; see equation (3.13). Collective conformational changes can also be represented via reconstructing the conformations from the (normally distributed) latent variable ***y***_*m*_ using equation (3.14). Interestingly, RPCA provides a clear distinction between the directions (generalized eigenvectors) along which the fluctuation of the conformations increases upon the change (corresponding to *λ_i_* > 1) and the directions along which the fluctuation of the conformations decreases (corresponding to *λ_i_* < 1). Figure 5a shows the collective conformational changes around the average conformation of the bound state along the 33 eigenvectors with the highest generalized eigenvalues (*λ_i_* > 10) which represent the enhanced motions of the wild-type HIV-1 protease upon binding Tipranavir. Figure 5b, on the other hand, shows the collective conformational changes around the average conformation of the free state along the 33 eigenvectors with the smallest generalized eigenvalues (*λ_i_* < 0.009)) which, in turn, represent the most strongly restricted motions of the wild-type HIV-1 protease upon binding Tipranavir. It should be stressed that the importance of the local conformational changes (e.g. at residues) cannot be inferred from the collective conformational changes. In other words, a large fluctuation in a site, when presenting the collective conformational changes, does not imply larger thermodynamic importance than the associated less strongly fluctuating sites. Alternatively, hotspot analysis from RPCA in equation (3.15) can be used to rank the thermodynamic contribution of the local conformational changes (conformational hot spots) of the biological building blocks (residues) to the association process and the importance of the cooperative (correlation) interactions between them. Figure 6a shows a matrix plot representation (interaction map) of the contributions of the residues to the divergence. In Figure 5b, the relative local residue contributions to the divergence are mapped on the structure of the protein and the importance of the conformational changes is indicated by radii of varying size in the cartoon representation, in addition to using the color code. Interestingly, most of the marked hotspots are residues known to affect the binding affinity upon mutating them. Examples of the defined conformational hot spots are the residues in the active pocket (e.g. D25, V82) and the residues of the flap region (e.g. I50, I54). Figure 7a shows an analysis of the conformational hotspots from RPCA of the binding of a ligand (Saquinavir) to the HIV-1 protease mutant with a resistance-related mutation (I50V) which is located on the flap region outside the binding pocket. The conformational hotspots of the mutant are located at the flap region around the mutation V50. The same flap region does not show important conformational changes when applying the same analysis to the association of Saquinavir to the wild-type (I50) in Figure 7b.

**Figure 5.**
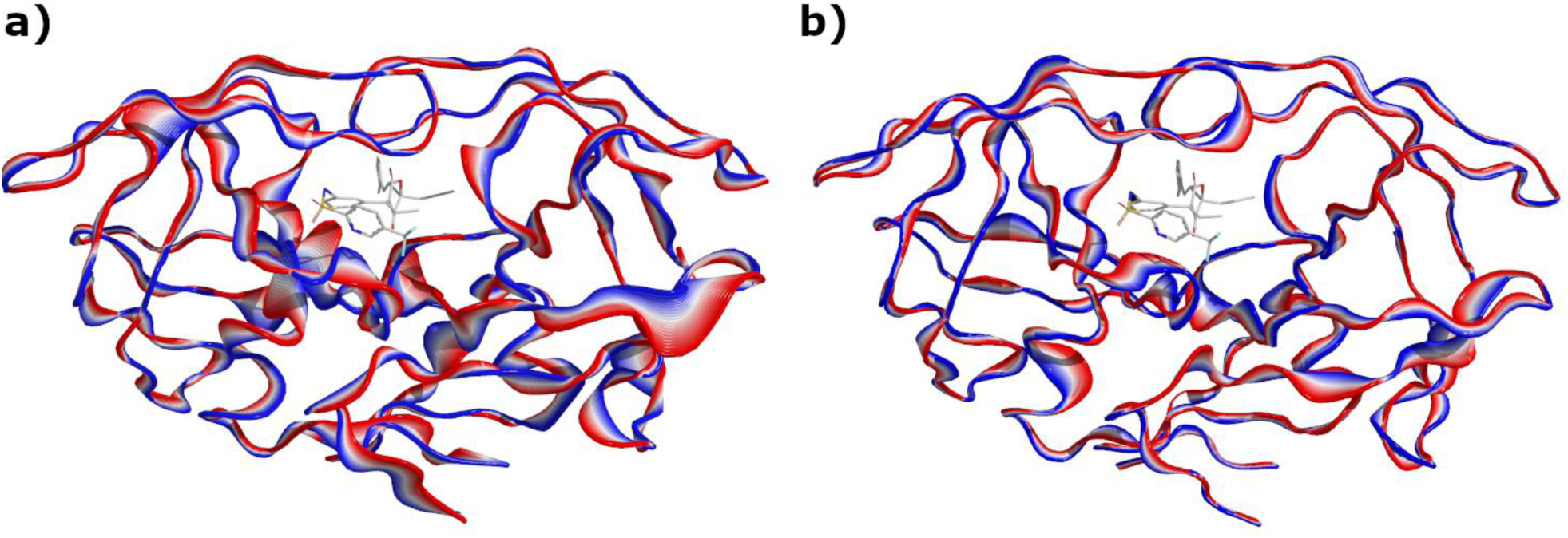
Representation of the enhanced and restricted conformational fluctuations of HIV-1 protease upon binding the inhibitor Tipranavir. Conformations around the average conformation are reconstructed from the latent variable after interpolation around its average along selected generalized eigenvectors; see equation (3.14). **a)** Enhanced conformational fluctuations around the average structure of the bound state along the 33 eigenvectors with the highest generalized eigenvalues (*λ_i_* > 10). These conformational fluctuations increase the affinity of binding by optimizing the local conformations in the ligand-receptor interface. **b)** Conformational fluctuations around the average structure of the free state along the 33 eigenvectors with the smallest generalized eigenvalues. These latter fluctuations are highly restricted upon the association (*λ_i_* < 0.009) because they decrease the affinity of binding via adverse local movements in the binding pocket and opening of the flap regions; see the supplementary movies. The cartoon representation is generated using Pymol (Schrödinger, LLC, 2018). The conformation of the ligand is taken from the experimental structure. Movies of these conformational changes are presented in the Supplementary Material S6.

**Figure 6.**
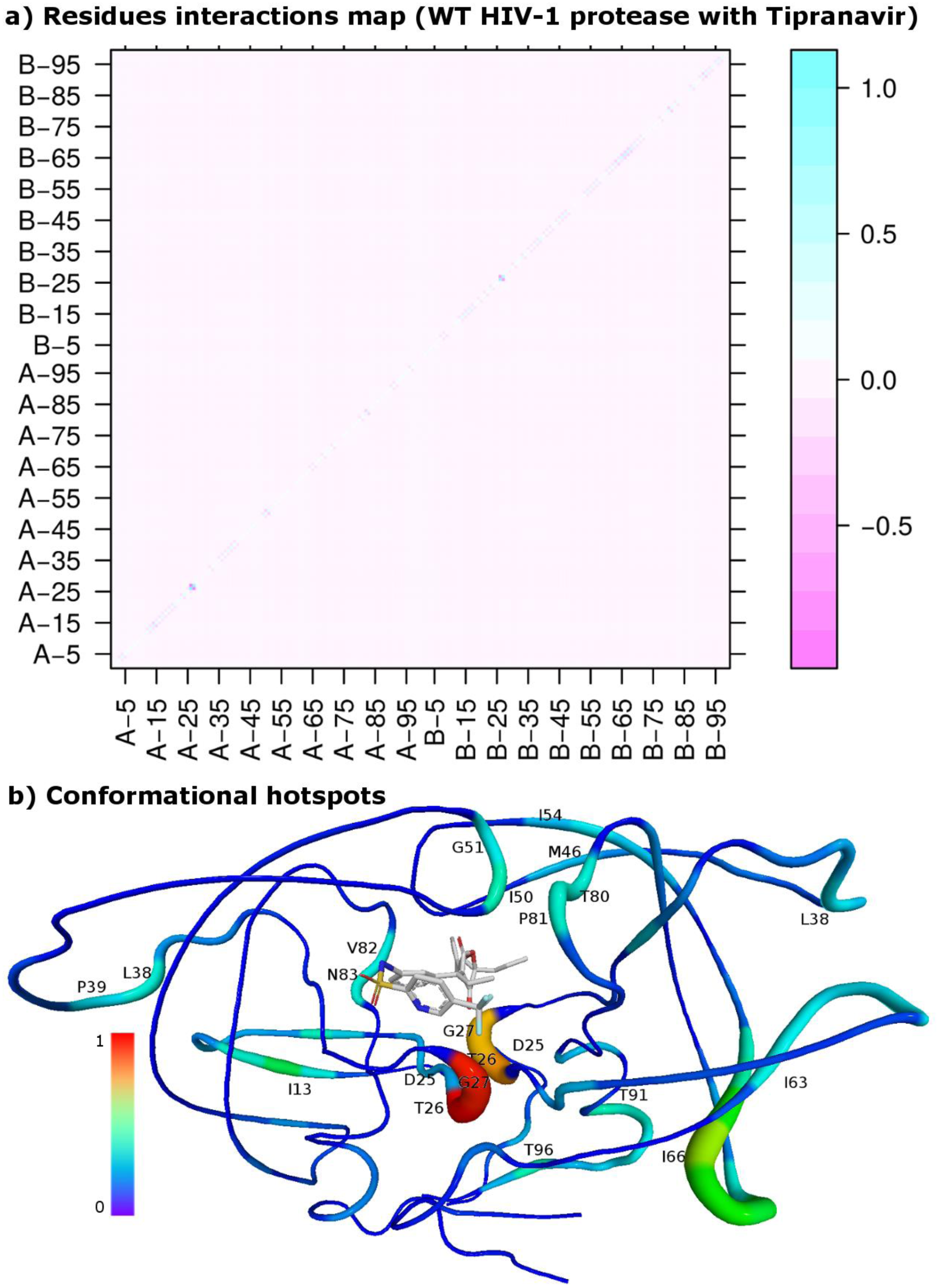
**a) Conformational interaction map** between the residues of HIV-1 protease upon binding its inhibitor Tipranavir. The contributions are concentrated around the diagonal elements indicating the important local conformational changes and the propagation of the conformational changes through the neighbor residues. **b) Conformational hotspots** mapped on the structure of the protein. The radius and the color of the cartoon indicate the importance of the conformational changes. The importance is normalized relative to the highest value that is set to 1.

**Figure 7.**
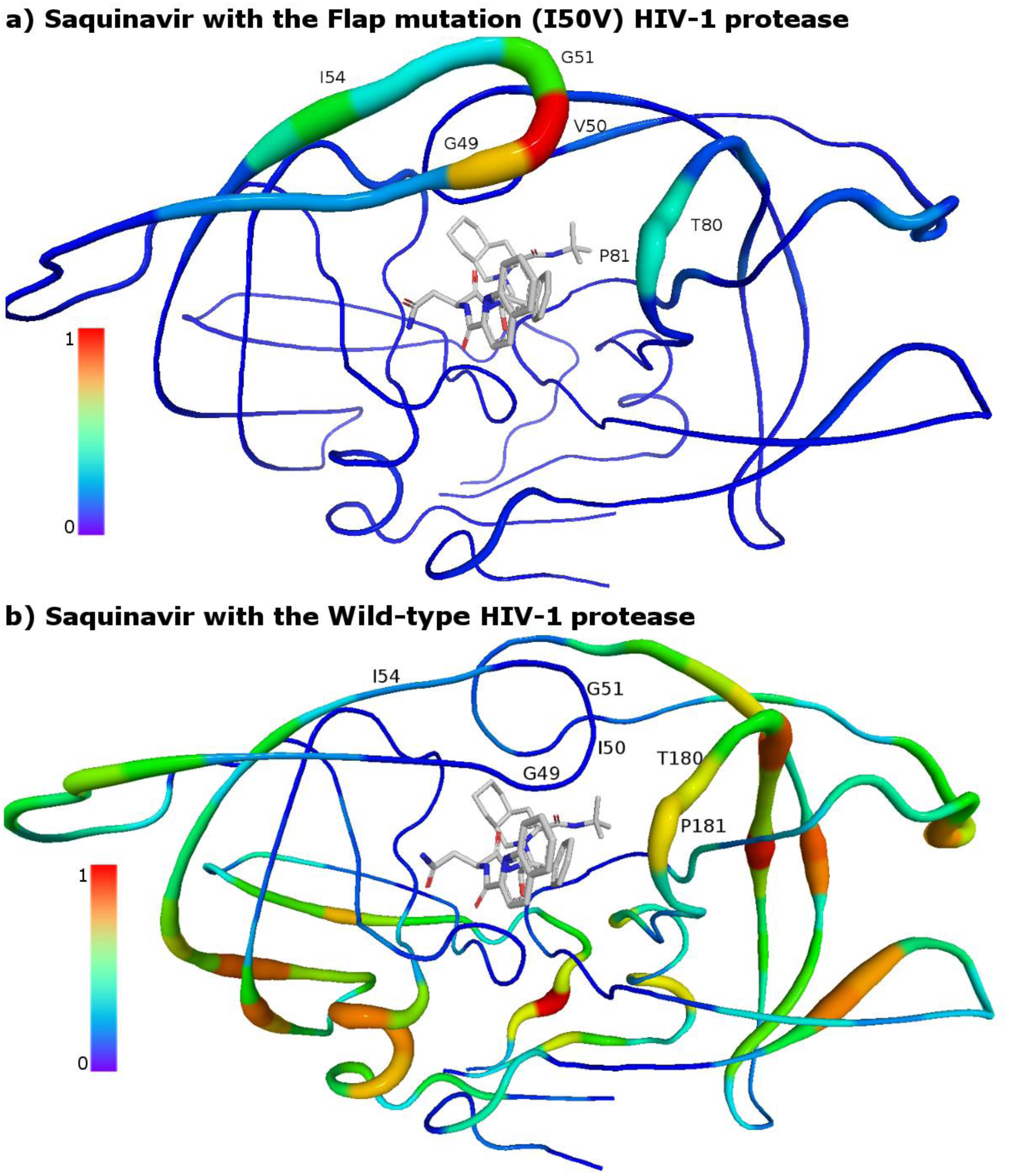
Conformational hotspots from RPCA analysis recognize the importance of the resistance-related mutation (I50V) of HIV-1 protease when bound to Saquinavir. The conformational hotspots of the mutant (a) are located at the flap region around the mutation V50. The corresponding flap residues of the wild-type (I50) in figure (b) do not show energetically important (expensive) conformational changes. The radius and the color of the cartoon indicate the importance of the conformational changes. The importance is normalized relative to the highest value.

### Technical aspects of using RPCA of molecular conformational changes

#### Optimal superposition of conformations of the ensembles

The first step in analyzing an ensemble of conformations is removing the six external rotational and translational degrees of freedom via superimposing the conformations. Although the internal coordinates are not altered due to these similarity transformations (rotations and translations), the average conformation and the covariance matrix are highly affected by the way of superimposing the conformations to remove these external degrees of freedom. Traditionally, the conformations are superimposed on an arbitrary reference structure (e.g. the starting structure of the MD simulation). However, the resulting ensemble and the average structure are dependent on the reference structure. The solution of this problem is well known in statistical shape analysis as the Generalized Procrustes Analysis (Dryden and Mardia, 2016; Gapsys and de Groot, 2013) (GPA). GPA returns a compact ensemble of the conformations via minimizing the sum of the Euclidean distances between the conformations (which is equal to their Euclidean distances to the average conformation). GPA is performed via an iterative algorithm having two steps: (1) compute the average structure, (2) superimpose the conformations on the new average structure. Repeating these steps ensures the convergence of the average structure. Fortunately, few iterations are usually enough for convergence.

#### Superposition of two ensembles

The Mahalanobis distance term 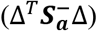 of the KL divergence in equation (3.9) is affected by the fashion in which we superimpose the average conformations (***μ***_*b*_, ***μ***_*a*_). However, superimposition via minimizing the Euclidean distance may overestimate the KL divergence via an artificial contribution to the Mahalanobis distance 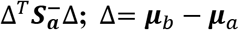. Therefore, superimposition of the average conformations should aim at minimization of the Mahalanobis distance. In contrast to the minimization of the Euclidean distance, there is no analytical solution known for minimizing the Mahalanobis distance. This type of nonlinear minimization is known as the Covariance Weighted Procrustes Analysis (Brignell et al., 2016) (CWPA) and numerical methods can be used to find the optimal superposition (rotations and translations) for minimizing the quadratic term 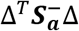.

#### Steps for performing a RPCA analysis of two molecular states

(1) GPA fitting of the conformations sampled in the simulation of the first state. The covariance matrix of this state is also computed at this step. A successful superimposition of the ensemble will lead to a singular covariance matrix with at least six eigenvalues of zero value accounting for removing the external degrees of freedom.
(2) GPA fitting of the conformations sampled in the simulation of the second state to obtain the average conformation.
(3) Covariance weighted fitting of the average conformation of the second state on the average conformation of the first state via minimizing their Mahalanobis distance. This unconstrained nonlinear optimization is numerically performed using the line-search algorithm and the BFGS factored method to update the Hessian (Dennis and Schnabel, 1996, p. 356).
(4) The new average conformation of the second state is used as a reference to refit the conformations of the second state and to compute the covariance matrix of the second state.
(5) Simultaneous diagonalization of the covariance matrices is performed. Optionally, the sub-spacing optimal algorithm can be used.
(6) KL divergences of the relative principal components are computed and the components are reordered based on their scores (KL divergences).

We have developed efficient computational tools to perform these steps. The tools are written in the C programming language and the numerical linear algebra operations are performed using the BLAS and LAPACK routines (Anderson et al., 1999). The limit of the zero value of the eigenvalues is defined using the machine precision multiplied by the largest eigenvalue. The covariance-weighted superimposition is performed using the nonlinear minimization algorithm by Dennis and Schnabel (Dennis and Schnabel, 1996, p. 356).

## 5 Conclusion

Here, we introduced the RPCA method, which extracts the relevant principal components describing the change between two macroscopic states of a dynamic system represented by two data sets. The definition of the macroscopic change of a dynamic system and its quantification are based on previous work where we derived a generalized quantification of the change of a physical system based on statistical mechanics. We presented use cases for conformational changes taking place upon ligand binding to HIV-1 protease that clearly illustrate the power of RPCA to characterize the relevant changes between two ensembles in a high-dimensional space. Moreover, software solutions were introduced to ensure the removal of the similarity transformations (rotations and translations) via superposing the conformations using GPA and CWPA. These procedures may also be beneficial for preparing the conformations for other analysis methods.

Although RPCA is currently limited to handling only continuous variables and two macroscopic states, the introduced framework for quantifying changes of dynamic systems using the exponential family of distributions is flexible regarding the nature of probability distributions and the nature of the microscopic variables (continuous and categorical). Therefore, the presented theoretical formalism opens the door for developing improved new methods for mining the factors underlying changes in dynamic systems in the directions of (i) handling both continuous and categorical data (e.g. the effect of sequence changes (mutations) on the binding affinity) and of (ii) handling multiple macroscopic states (e.g. study the binding of a series of ligands to a receptor).

